# Animal models in psychedelic research – Tripping over translation

**DOI:** 10.64898/2026.01.14.699469

**Authors:** Muad Y. Abd El Hay, Ana Cukić, Marieke L. Schölvinck, Martha N. Havenith

## Abstract

Psychedelic substances show promise for treating psychiatric disorders including depression, anxiety, and PTSD, but the neurobiological mechanisms underlying their therapeutic effects remain poorly understood. Preclinical findings have so far been inconsistent, limiting mechanistic insights. We argue this translational gap is exacerbated by systematic disconnects between conditions thought to shape therapeutic outcomes in humans and those employed in animal research. In this review, we analyse 266 rodent psychedelic studies published between 2014 and 2026 in order to generate a systematic audit of experimental conditions and behavioural assays currently applied in the field. Specifically, we first assessed adherence to welfare practices paralleling “set and setting” factors critical to human therapy. We found that most studies contained stress-inducing factors: only a minority reported active-phase testing (14%), environmental enrichment (7%), or refined handling protocols (21%), while drug administration almost universally relied on stress-inducing, restraint-requiring methods. Second, we evaluated the capacity of behavioural assays to capture experiential dynamics, revealing that most studies rely on brief, constrained testing with isolated behavioural markers that fail to capture the multidimensional, temporally evolving nature of psychedelic states. Beyond these methodological gaps, we identify a third critical factor largely absent from preclinical research: individual differences and social context. Most animal studies use group-level, cross-sectional designs that cannot characterize heterogeneous treatment responses or social mechanisms implicated in therapeutic change. We propose that improving translation requires a fundamental reorientation: from brief testing of stressed, isolated animals toward longitudinal tracking of individuals in enriched social environments using comprehensive behavioural characterization. We end by offering practical advice on how to implement some of these considerations in animal psychedelic research.

## Introduction

Mental disorders such as depression and anxiety impact the lives of an ever-growing number of people, often in persistent and debilitating ways. Globally, depression affects a staggering 280 million people (1). Yet despite decades of research and pharmaceutical development, nearly 40% of all patients with depression fail to respond to available treatments (2). This situation necessitates new treatment approaches to mental disorders that go beyond conventional drug development paradigms.

Among the most promising new alternatives, psychedelic substances have emerged as potential therapeutic tools. Substances like psilocybin, LSD, and MDMA, some of which have been used in traditional practices for centuries, are the subject of 488 clinical trials to date – tendency rising (3). These trials show that psilocybin has therapeutic efficacy comparable to established antidepressants, with a notably favourable side effect profile (4). Importantly, these effects extend even to patients that were deemed ‘treatment-resistant’ (5), and they apply across a wide range of prominent mental disorders including anxiety, post-traumatic stress disorder (PTSD), and even addiction (6–8) – a broad applicability that is uncommon for classical treatments. Moreover, therapeutic changes are typically achieved in only one or two dosing sessions, yet produce long-lasting improvements (4).

Together, these properties position psychedelic substances as a potential ‘therapeutic multitool’, addressing a wide variety of mental ailments, often in a one-shot treatment. What processes allow psychedelics to trigger such substantial changes in neuronal processing in such a short time, and across vastly different mental health conditions? At a global level, psychedelic substances have been shown to disrupt ordinary brain organization through widespread desynchronization, decreasing activity in the default mode network while dissolving boundaries between typically segregated brain regions (9–11). But how are such global shifts in neuronal circuit communication produced at the circuit level, and how do they in turn translate to lasting therapeutic changes? To date, these questions remain largely unresolved, and due to their non-invasive nature, human brain recordings lack the necessary spatial or temporal resolution to tackle them directly. As such, we must study the neuronal circuit mechanisms of psychedelics in animal models. Given their potential for detailed circuit-level recordings and controlled behavioural testing, animal models are essential for drawing mechanistic links between acute shifts in neuronal circuitry triggered by psychedelics, and their long-term effects. However, psychedelic research in animal models has so far mainly succeeded in establishing receptor signaling pathways and acute neuroplasticity markers, whereas the factors thought to shape therapeutic outcomes in humans—including the proposed role of context, the predictive significance of acute experiential quality, and the substantial heterogeneity in treatment response—have largely remained unaddressed.

In this analysis of the field, we argue that the successful translation of psychedelic research from animal models to human psychedelic therapy hinges on three factors that are well established in clinical research, but have been largely overlooked in preclinical studies: (1) The importance of set and setting (the psychological mindset and environmental context in which psychedelics are administered), (2) The central role of subjective experiences for acute and long-term effects, and (3) The significance of inter-individual differences for therapy outcomes. We will examine how current animal models address each of these factors, and propose methodological advances that could improve translation between preclinical findings and clinical outcomes.

### Set and Setting

The concept of “set and setting” has been recognized as a critical determinant of subjective experiences and therapeutic outcomes in human psychedelic research. “Set” refers to the individual’s mindset, expectations and emotional state, while “setting” refers to the physical and social environment in which the psychedelic experience occurs. A strong therapeutic alliance and a mental state of surrender are associated with more positive therapeutic outcomes, possibly by encouraging deeper psychedelic experiences (12–14), while feelings of preoccupation prior to a psychedelic session and greater anxiety during the experience predict less favourable clinical outcomes (14, 15). Self-reports from recreational users similarly show a strong correlation between feelings of safety during the experience and positive long-term effects (16). While the causal contributions of individual contextual factors are only now being dissected in randomised trials (17, 18), the perceived importance of context in psychedelic therapy has already motivated the development of standardized reporting guidelines (19) as well as recommendations aimed at increasing the likelihood of desirable outcomes and reducing adverse effects through set and setting factors (20).

Given the central role of environmental, psychological and experiential factors in human psychedelic therapy, we examined whether preclinical studies account for analogous considerations in animal models.

Rodents are the most commonly used animal models in translational research on psychedelics. Initial findings suggest that psychedelic episodes in rodents show context effects and altered sensory processing, closely echoing similar processes in humans (21–24). This suggests that “set and setting” factors are also crucial in animal models in order to capture therapeutically relevant psychedelic processes. We therefore systematically examined how psychedelic research has implemented key welfare practices by analysing the housing, handling, and experimental conditions in 266 rodent studies published from 2014–2026 (Figure 1 and Supplementary Fig. S2).

**Fig. 1.**
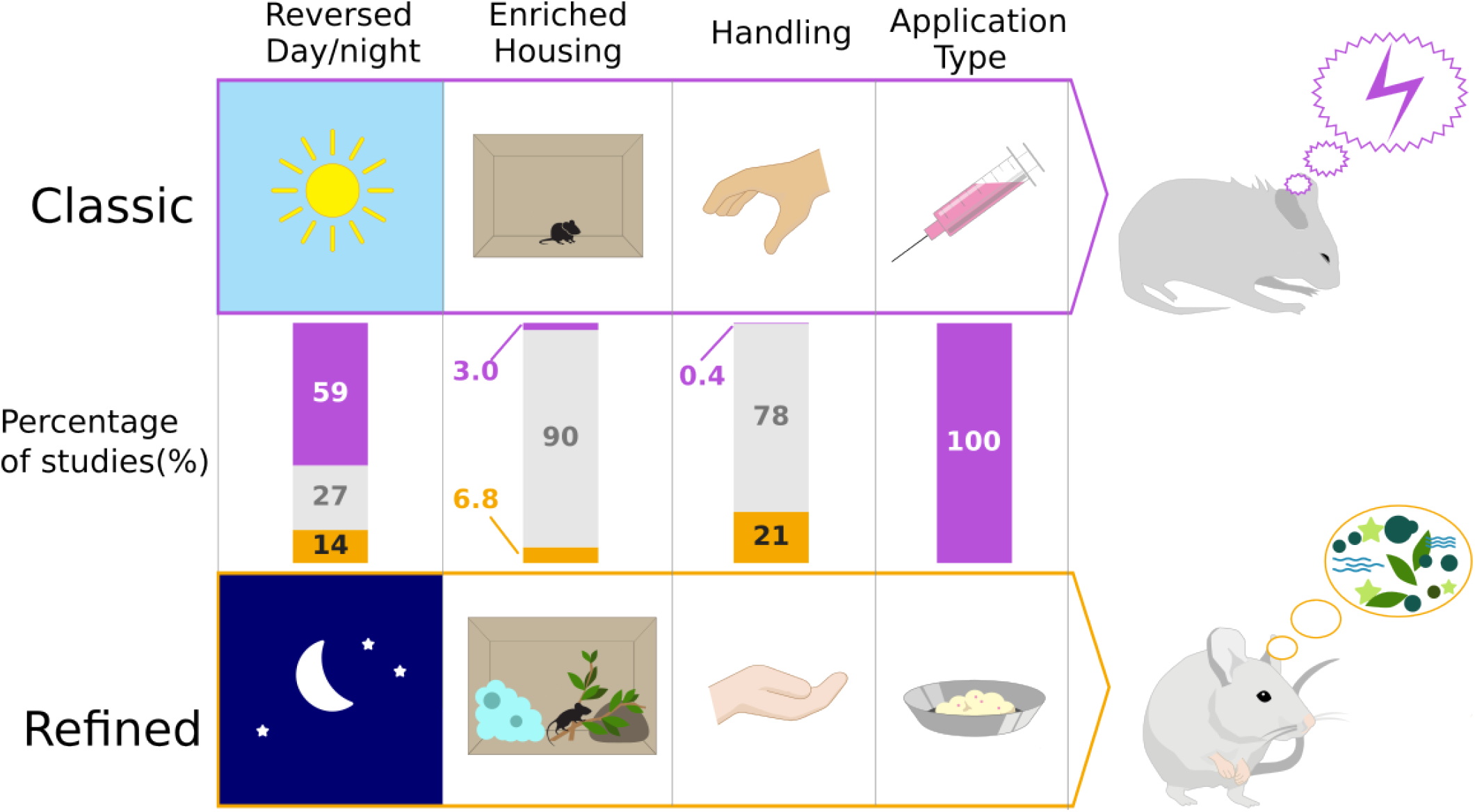
Comparison of classic versus refined experimental practices in psychedelic animal research. Based on our systematic review of 266 rodent studies (2014–2026), the figure contrasts prevalent ‘Classic’ practices (purple) with recommended ‘Refined’ practices (orange) across four key parameters: circadian alignment (reversed day/night cycle), environmental enrichment and communal housing, handling and habituation protocols, and drug application method. Each bar splits the corpus into three segments: studies that explicitly used the refined practice (orange), studies that did not report the practice (grey), and studies that explicitly used the classic practice (purple). Classic practices create stressed experimental conditions, while refined practices promote natural behavioural states (depicted by calm mouse in an enriched setting).

#### Circadian alignment

Consistent with research in humans, research in rodents has shown that the circadian phase of testing can substantially influence behavioural outcomes. Locomotor activity is consistently higher during the dark (active) phase, while effects on social behaviour and cognitive performance are more task-dependent (25–27). Rodents are predominantly nocturnal animals, with peak activity occurring during the dark phase of the light/dark cycle. Yet of the 266 psychedelic studies we analysed (Figure 1), only 37 explicitly reported housing rodents under a reversed light/dark cycle to enable active-phase testing, while 158 studies adhered to a standard cycle, and 71 did not report this information. This indicates that the majority of psychedelic experiments in rodents were conducted during their inactive (sleep) phase. This mismatch likely contributes to poor translation of preclinical findings by altering the animals’ cognitive, emotional, and physiological state during psychedelic exposure. Notably, these phase-dependent effects extend to drug responses themselves: antidepressants from different classes show distinct chronopharmacological profiles in rodent assays, with peak efficacy depending on dosing time (28), and the circadian clock has been implicated in the action of rapid-acting antidepressants such as ketamine (29).

#### Environmental enrichment and communal housing

Environmental enrichment provides animals with complex physical, sensory, and social stimulation that better meets species-specific needs, thereby reducing stereotyped behaviours, improving welfare, and allowing animals to express more naturalistic behavioural repertoires. Enrichment has been shown to reveal richer phenotypes without increasing data variability (30–33), while at the same time reducing morbidity and mortality, compared to standard laboratory housing (34). Notably, environmental enrichment alters psychedelic-induced brain activity (35), and is a prerequisite for cognitive improvements following LSD exposure in ageing rats (36). The same pattern extends beyond psychedelics: housing condition modulates the efficacy of SSRIs and SNRIs (37–39), flips the direction of ketamine’s effect in the forced swim test between stressed and unstressed mice (40, 41), and blunts the rewarding effects of cocaine, fentanyl, and alcohol (42–45). Home cage monitoring in enriched environments can thus capture temporal patterns, social behaviours, and spontaneous activities of animals that improve translational relevance (46, 47). In our analysis, only 18 of 266 studies reported providing enriched home cages, while 8 studies explicitly stated no enrichment and 240 did not report this information (Figure 1).

In addition to physical enrichment, whether rodents are housed individually or in groups has profound effects on their well-being. Social isolation is a significant stressor that produces anxiety- and depression-like behaviour, cognitive deficits including impaired memory and social cognition, and altered immune and metabolic function (34, 48, 49). As such, individual housing in psychedelic experiments is highly likely to create confounds that may mask or distort experimental effects. Group housing can reduce such adverse effects by providing species-appropriate social contact, though the magnitude of isolation effects varies with strain, sex, and duration (50). Isolation also directly reshapes drug responses: isolation rearing potentiates psychostimulants (51) and ethanol intake (52), blunts fluoxetine efficacy in genetically vulnerable rats (53), and the drug status of cage-mates modulates morphine dependence and conditioned place preference (54). In our analysis, 157 studies housed animals in groups, 46 reported individual housing, and 63 did not specify housing conditions. Together, these findings indicate that solitary or unspecified housing conditions in a substantial share of studies may curtail the therapeutic mechanisms of psychedelics.

#### Handling and habituation protocols

Handling technique and the extent of habituation to human contact profoundly im-pact rodent stress levels and behaviour. For example, conventional tail handling induces an anxiety-like state that changes how animals perceive and respond to rewards (55), while proper handling protocols markedly reduce stress responses and variability in behavioural tests (56–58). Handling is not merely a behavioural confound: gentle, habituated handling raises peak plasma drug concentrations by*∼* 30% relative to tail lifting in a controlled pharmacokinetic study (59), and the sex of the human experimenter modulates ketamine’s antidepressant action in mice (60). In our analysis, handling was reported in only 57 studies, while 1 study explicitly stated that no specific handling was used, and 208 did not provide this information (Figure 1).

Similarly, familiarization with the test arena aims to reduce novelty-induced stress and anxiety, allowing researchers to measure experimental effects rather than stress confounds. Brief arena familiarization has been shown to increase stimulus investigation and improve habituation–dishabituation performance in mice, with effects most pronounced in more anxious animals (57). Repeated exposure to test environments reduces novelty-induced anxiety and improves the reliability of exploratory behaviour measures (61), thereby enhancing statistical power and reproducibility. In our analysis, familiarization to the experimental setup was included in 153 studies, while 39 did not incorporate a habituation period, and 74 studies did not mention whether habituation was performed (Figure 1). Given that environmental context and concurrent stress shape both brain-wide neural activity and acute and post-acute responses to psychedelics in rodents (35, 62), a lack of habituation may introduce substantial variability by conflating novelty-induced stress with psychedelic effects.

#### Drug administration (Figure 1)

The route and method of drug administration represent an often overlooked welfare consideration that directly impacts the stress levels with which animals enter psychedelic experiences. Restraint, which is required for intraperitoneal injection, subcutaneous injection, and oral gavage, is itself a standard method to experimentally induce stress in rodents (63, 64). Evidence demonstrates that these routine administration procedures trigger substantial physiological stress responses, including elevated heart rate, increased body temperature, and raised corticosterone levels that can persist for several hours (65–67). In contrast, voluntary oral administration methods using palatable vehicles have been developed to minimize such stress, producing lower corticosterone levels and more stable cardiovascular responses while preserving target engagement (67–69). Also, administration route can itself alter drug efficacy: in mice undergoing chronic unpredictable stress, fluoxetine produced larger antidepressant effects when given intraperitoneally than by intragastric gavage, with gavage itself inducing a stress-like behavioural phenotype (70).

In our analysis, the vast majority of psychedelic studies relied on injection methods (Figure 1), with intraperitoneal (i.p.) administration accounting for 69% of the per-study route weight, followed by subcutaneous (s.c.) at 15%, and oral gavage (p.o.) at 10% (Supplementary Fig. S3). Notably, despite the availability of voluntary oral administration methods that could minimize handling-related stress and better approximate a human clinical context, this approach appears to be entirely absent from the psychedelic literature. The predominant use of stressful administration methods raises an important question: can we expect to faithfully model the therapeutic effects of psychedelics when animals experience drug onset in a state of acute anxiety – a context that, in clinical reports, has been associated with poorer outcomes in human patients (14, 15)?

#### Choice of animal model

Whether and how these welfare practices shape experimental outcomes depends in part on which species is being studied. Mice and rats differ markedly in default stress profiles and behavioural repertoires: rats show larger home ranges, slower-paced exploratory profiles, and richer learned-behaviour repertoires than mice (71–74). The two species also occupy distinct stress profiles: mice engage in risk-assessment behaviours on initial exposure to threatening stimuli that rats do not, while rats produce alarm vocalisations absent in mice (75), and meta-analyses of maternal separation find that the same early-life stressor durably increases adult anxiety-like behaviour in rats but leaves it largely unchanged in mice (76). Welfare and procedural standards that are appropriate for one species may therefore not be appropriate for the other. In our review, 157 of 266 studies (59%) used mice, 99 (37%) used rats, and 10 (4%) used both, yet across all four welfare practices analysed, mouse and rat studies showed closely matched profiles, differing by no more than 4 percentage points: reversed light cycle was reported in 15% of mouse studies versus 13% of rat studies, enrichment in 8% versus 6%, group housing in 58% versus 62%, and refined handling in 19% versus 25%. This uniformity suggests that current welfare reporting is driven more by convention than by species-specific consideration of stress sensitivity.

Given that clinical psychedelic therapy is typically conducted within supportive, low-anxiety contexts thought to facilitate positive therapeutic outcomes, the widespread neglect of analogous welfare practices in preclinical animal research (including environmental enrichment, stress reduction, active-phase testing, and careful handling) may compromise our ability to model and understand the neurobiological mechanisms underlying the therapeutic effects of psychedelics.

One recent clue that the therapeutic mechanisms of psychedelics may indeed be somewhat ‘lost in translation’ comes from a large-scale, multi-institutional preprint testing the effects of psilocybin using approximately 200 mice per experiment across five independent laboratories (77). Despite strong statistical power and transparent reporting, the study failed to find replicable therapeutic effects of psilocybin 24 hours post-administration. Specifically, psilocybin neither reduced behavioural markers of anxiety or depression, nor did it boost fear extinction or social reward learning. The only replicable effects were acute, including decreased fear expression and increases in anxiety and avoidance behaviours during drug administration.

While these results may point towards a low clinical utility of psilocybin and/or a strong divergence in the effects of psilocybin on humans and rodents, another possibility is that the observed effects were strongly shaped by experimental context: Like most studies surveyed in our analysis, Lu et al. (77)did not include environmental enrichment, handled animals by scruffing, administered psilocybin via intraperitoneal injection, and conducted 80% of their experiments during the animals’ inactive circadian phase.

Together, these aversive context factors may have had a sufficiently strong impact to counteract therapeutic mechanisms that might be engaged by psilocybin under different circumstances. This possibility also tallies with the study’s finding that animals’ anxiety increased consistently during drug administration — an effect one would expect to find in “bad trips” in a human context. These possible interpretations underscore the notion that, particularly in psychedelic research, welfare practices may not be peripheral considerations, but rather experimental parameters that could critically shape the success of preclinical studies as a translatable model of therapeutic processes triggered by psychedelics in humans.

#### Dose equivalence across species

A further translational concern, tied to the question of behavioural readouts, is dose equivalence: at what dose does a rodent enter a state comparable to the human psychedelic experience? Two indirect measures are used to triangulate this: head-twitch response (HTR) intensity in rodents and 5-HT_2*A*_ receptor occupancy measured by PET. For psilocybin these converge reasonably well (78, 79). Mouse PET data place peak HTR intensity at 44–62% cortical 5-HT_2*A*_ occupancy (78), overlapping the 40– 70% window that produces psychedelic and therapeutic effects in humans at 10–30 mg oral psilocybin (79, 80). Translated to rodents this corresponds to approximately 0.6–2 mg/kg in mice (78) and 1–4 mg/kg in rats (81), and the doses used across the reviewed psilocybin studies fall largely within this range in both species (Supplementary Fig. S4). For LSD, no *in vivo* 5-HT_2*A*_ occupancy curves have been reported in any species; for DMT, a recent rat PET study (co-administered with the monoamine oxidase inhibitor harmine) found no measurable 5-HT_2*A*_ occupancy even at brain DMT levels up to 11.3 µM (82). Even where HTR and occupancy converge, however, they remain pharmacological proxies and cannot tell us whether the animal is in a phenomenologically comparable state. Establishing this calls for direct tracking of the animal’s behavioural and neural state over time, an approach discussed below.

### Tracking experiential dynamics

In humans, psychedelic substances induce a multitude of subjective experiences ranging from simple sensory alterations such as visual hallucinations and audiovisual synaesthesia, to complex cognitive changes such as disembodiment or impaired cognitive control. Perhaps most significantly, psychedelics can induce strong emotional responses such as feelings of catharsis and awe, as well as spiritual or mystical experiences, including feelings of unity and transcendence of space and time (83). Human studies have revealed that the subjective quality of acute psychedelic states predicts longterm therapeutic outcomes (84–86). Specifically, participants that report experiencing mystical or meaningful experiences during their sessions often achieve better long-term outcomes such as a reduction in depressive symptoms (15, 87), while participants who report aversive or anxious experiences may derive less benefit (88).

This raises a challenge for translational research: we cannot access the subjective experiences of animals through verbal report, but this does not imply that animals lack rich experiential states during psychedelic exposure. On the contrary, behavioural observations across species suggest that their subjective experiences may be noticeably altered. Cats under the influence of psychedelics grasp at the air with their paws, visually track objects not present in the environment, and pounce toward empty spaces (89, 90). Similarly, rhesus monkeys show “tracking” behaviour of invisible objects after hallucinogen administration (91). Most compellingly, recent work has directly demonstrated parallel visual distortions in humans and rats exposed to psilocybin using an identical behavioural paradigm (21). These converging observations indicate that animals are not merely exhibiting pharmacologically induced motor dysfunctions, but rather appear to respond to altered perceptual states that shape how they interact with their environment.

The problem, therefore, is not that animals necessarily lack subjective experiences during psychedelic exposure, but rather that current animal studies have rarely characterized these experiential dynamics. Most preclinical psychedelic research relies on isolated behavioural markers such as head twitches, total amount of locomotion, or brief snapshots of social interaction—measured at single time points or in highly constrained test conditions. This reductionist approach, while maximizing experimental control, may compromise translational validity (92–95). The head twitch response exemplifies this principle: It is reasonably straightforward to quantify, and shows remarkable predictive validity for identifying psychedelic compounds (96). At the same time, its face validity compared to human phenomenology is more limited, and on its own it may tell us relatively little about the quality of the animal’s experiential state, for instance whether perceptual changes it may be experiencing are felt as aversive or pleasant. The current scarcity of standardized approaches for characterizing acute experiential states in more detail may also be one reason why it can be challenging to determine why the same compound is sometimes reported to produce anxiolytic effects and in other cases anxiogenic effects (62, 97). Such conflicting outcomes might be explained by contextual factors that alter the animal’s overall experiential state, but we can only test this hypothesis fully if metrics are available to differentiate qualitatively distinct experiential states.

To address this gap, we propose that animal research must move beyond simple behavioural markers toward comprehensive characterisation of experiential dynamics during psychedelic exposure. While we cannot directly measure subjective experience in animals, comprehensively tracking a multitude of behavioural components — such as altered sensory processing, changes in exploration-avoidance balance, social behaviour modifications, and temporal patterns of arousal and rest — allows us to characterize what constitutes a “psychedelic state” in animals and how different qualities of this state might predict downstream outcomes (93). Following the framework proposed by Shemesh et al. (92) for improving translational validity in psychiatric research, we identify three features that must be considered to adequately capture psychedelic-induced experiential dynamics in animal models: behavioural complexity, environmental diversity and temporal dynamics (Figure 2).

**Fig. 2.**
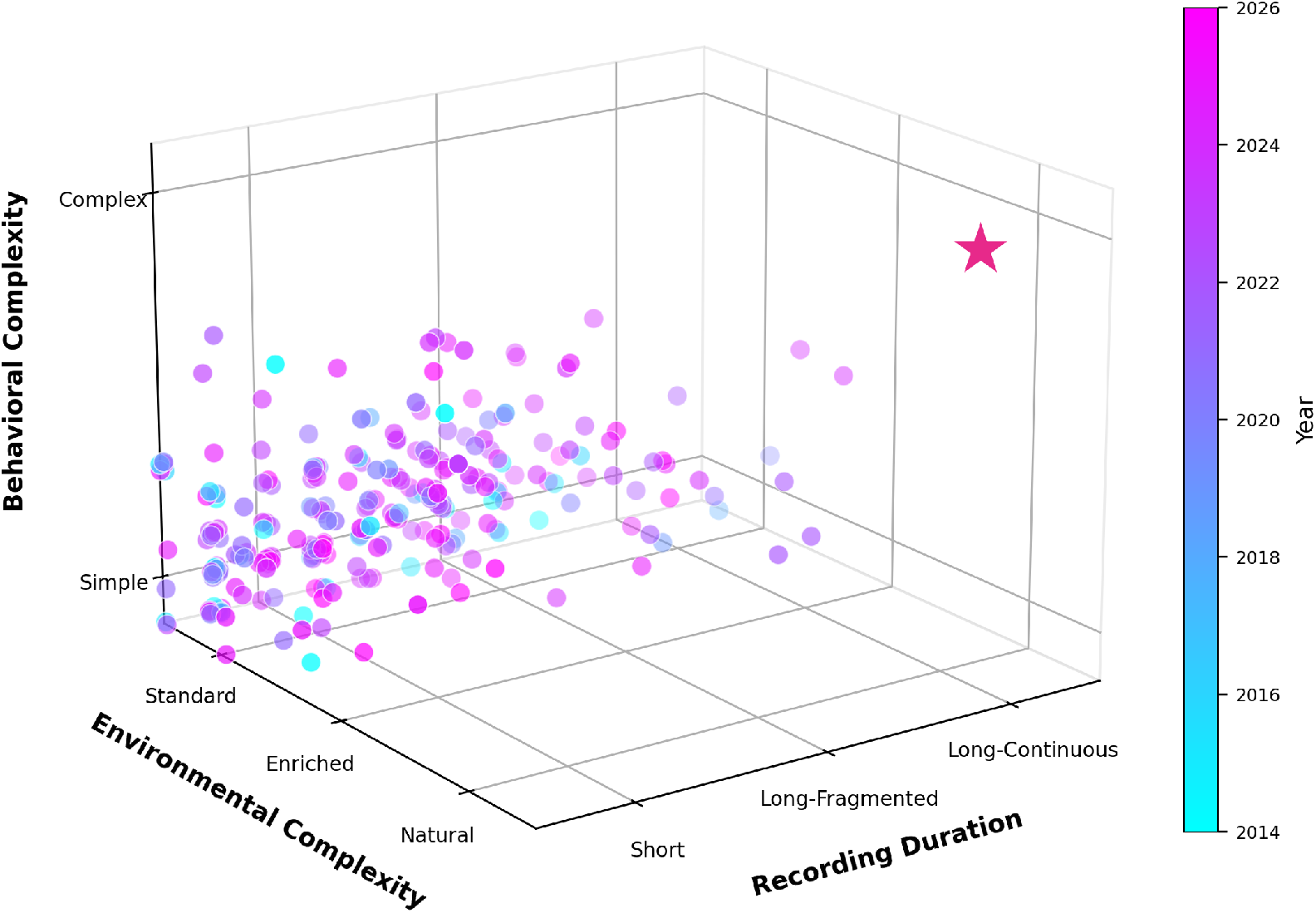
Mapping behavioural assays in rodent psychedelic research. Each point represents one of 266 reviewed studies, positioned in three-dimensional space according to recording duration (x-axis: from brief single-session tests to extended continuous monitoring), environmental complexity (y-axis: from barren test apparatus to semi-naturalistic habitats), and behavioural complexity (z-axis: from single metrics like head twitches to multidimensional behavioural characterization). For a detailed description of the calculation, see Materials and Methods. Point colour indicates publication year. The pink star marks an aspirational direction: comprehensive, continuous behavioural monitoring in naturalistic environments—the approach most likely to capture therapeutically relevant experiential dynamics.

#### Capturing behavioural complexity

Human subjective experiences under psychedelics are multidimensional, encompassing perceptual, emotional, cognitive, and social domains. Reducing animal behaviour to single metrics (e.g. number of head twitches) may miss the very aspects of behaviour that reflect translationally and therapeutically relevant experiential states. Advances in computational ethology—combining automated behavioural tracking, machine learning-based behavioural classification, and analysis of behavioural sequences—now enable comprehensive, multidimensional characterization of behaviour without requiring labour-intensive manual scoring (92, 93, 98). These computational approaches enable researchers to detect complex behaviours such as social approach and retreat, affiliative interactions, defensive responses, and their sequential organization. For example, rather than measuring total social interaction time, we might characterize the sequential structure of social behaviours (approach → investigation → withdrawal), the balance between affiliative and defensive responses, the flexibility versus predictability of behavioural transitions, accompanying vocalizations, and how these patterns evolve over the course of psychedelic exposure. Recent computational approaches further demonstrate the power of behavioural characterization in extracting the complexity of underlying experiential states. For instance, machine-learning analysis of facial expressions in mice can reverse-engineer emotional states from micro-expressions invisible to human observers (99), while ultrasonic vocalization analysis reveals that superficially similar behaviours can have opposite emotional valences (100). Such comprehensive behavioural characterization can capture measurable components of experience that may more faithfully reflect the experiential dynamics involved in the therapeutic effects of psychedelics in humans (93, 98).

#### Providing environmental diversity

In clinical settings, patients experience psychedelics in comfortable environments that allow for diverse behaviours including rest, movement, interaction with objects or music, and social connection with therapists. In contrast, most animal studies confine subjects to barren test environments that permit only a limited behavioural repertoire. This environmental impoverishment restricts the range of behaviours animals can express, potentially masking experiential dynamics that would be evident in more naturalistic settings (94, 95). For example, in a semi-naturalistic setting called the Visible Burrow System, rats clustered into ‘stress-responsive’ and ‘stress non-responsive’ phenotypes — a distinction that did not emerge in more confined settings (101). Similarly, psilocybin’s anxiolytic effects (quantified by the elevated plus maze task) only emerged when rats received repeated exposure to novel environments during the weeks following psilocybin administration (102). These findings suggest that beneficial versus detrimental effects of psychedelics— and the mechanisms that produce them — may only be tractable when animals are studied in sufficiently complex environments that enable a diverse behavioural repertoire (92, 98).

#### Recording temporal dynamics

Human psychedelic experiences unfold over hours, with distinct phases including onset, peak effects, and gradual return to baseline — each associated with different experiential qualities. In contrast, the majority of animal studies measure behaviour in brief snapshots, e.g. by applying a brief open-field test. Such measurements by nature cannot reflect how behavioural states evolve across the time course of psychedelic exposure, whether they transition between distinct phases, and how these in turn might predict later outcomes. Moreover, fragmenting behavioural readouts e.g. by moving animals between test apparatuses or repeatedly handling them for serial testing may itself disrupt the natural evolution of psychedelic states. Long, continuous, and noninvasive behavioural monitoring throughout the psychedelic experience is essential to characterize how behavioural states evolve spontaneously over time, identify critical transition points, and detect temporal patterns that may predict therapeutic long-term effects (92, 98).

Our analysis of current psychedelic research in rodents highlights that most studies fall short of these recommendations (Figure 2), and that this picture has not been improving: across the 12-year span of our corpus, none of the three behavioural-assay scoring dimensions shows a significant temporal trend (Supplementary Fig. S8). However, a small number of studies have begun to adopt more comprehensive approaches, demonstrating the feasibility and value of time-resolved recordings in enriched environments with improved behavioural characterization (35, 103). These pioneering efforts point toward a future where animal models can more faithfully capture the experiential dynamics that appear critical for therapeutic outcomes in humans.

### Understanding psychedelic action in context

#### Individual differences in treatment response

Clinical outcomes in psychedelic therapy are characterized by substantial individual variation: while some patients experience intense mystical experiences and lasting therapeutic benefits, others experience psychologically distressing sensations and minimal therapeutic responses or even adverse effects (88). For instance, in psilocybin trials for treatment-resistant depression, therapeutic success rates typically range from 50–70% (4, 5). Multiple sources of variation could contribute to individual differences in psychedelic response, including genetic factors (104, 105), psychological factors such as personality traits (14, 106, 107), and set and setting (14, 108). Understanding this heterogeneity represents a major challenge for bringing psychedelic therapy into safe and effective clinical practice, and preclinical research can play a crucial role in tackling it. This approach is particularly promising because even mice raised under standardized conditions show substantial individual variation in behaviour and brain structure, providing a translationally valid target for individualized outcome prediction (109, 110). While perhaps surprising, this variation is readily understood when considering that early-life contingencies can generate self-reinforcing individual differences, which in turn become amplified through competitive social feedback, pushing genetically identical individuals onto divergent life trajectories (111).

Despite their ubiquity, inter-individual differences have been largely neglected by psychedelic research in animals, which has predominantly focused on group-level effects rather than tracking individual animals over time. As such, individual differences in behavioural or neural responses are rarely systematically characterized, and when observed, they are typically attributed to measurement error rather than recognized as potentially reflecting the same response heterogeneity seen in humans. Sex is a further source of individual variation, and one whose role in psychedelic therapy is still being worked out. In humans, the clinical picture remains unclear: although estrogen–serotonin interactions provide a mechanistic basis for sex-dependent responses, controlled studies have so far not established robust sex differences in the subjective effects of psychedelics, and clinical trials continue to under-enrol and under-report female participants (112, 113). Animal studies, however, already reveal effects that this human work is not yet powered to resolve: female C57BL/6J mice show larger maximal head-twitch responses to DOI, LSD, and psilocybin at high doses despite comparable ED_50_ values (114, 115), psilocin produces sex-specific changes in central amygdala reactivity and threat behaviour (116), and ovarian hormones modulate 5-HT_2*A*_-mediated behavioural and circuit responses through direct estrogen–serotonin interactions (112). A recent meta-analysis of 124 studies further establishes that estrous-cycle phase robustly modulates anxiety-related behaviour (g*≈* 0.44 across rats and mice) (117). Animal models are therefore well placed to dissect exactly the sex-dependent mechanisms that human studies cannot yet resolve — yet our corpus shows the field is largely forgoing this opportunity. Of the 266 reviewed studies, 176 (66%) used male animals only, while just 76 (29%) included both sexes, 11 (4%) used female animals only, and 3 (1%) did not report sex (Supplementary Fig. S5). This is beginning to change: the share of studies including both sexes has risen sharply since 2022 (from less than 15% in 2014–2021 to roughly 40% in 2023– 2025), but male-only designs continue to dominate. Realising the translational potential of this question will require studies that include both sexes by design rather than as an exception, documenting estrous-cycle phase when females are included.

To further address individual traits that predict psychedelic response profiles, we must track how specific individuals behave from baseline through acute psychedelic exposure and into the weeks or months following treatment. Longitudinal designs in this vein allow us to ask central questions such as: How do baseline characteristics predict which animals will enter different acute experiential states, and how does this predict subsequent behavioural task performance? In human research, this approach has proven essential: studies tracking individuals across multiple time points have revealed that baseline brain network organization predicts treatment response (118), that changes in default mode network connectivity during acute psychedelic states correlate with symptom improvements weeks later (15, 119), and that individual differences in acute experiences mediate long-term outcomes (15). Yet in animal research, most studies employ between-subjects designs, preventing direct tracking of individual trajectories and thereby making it impossible to identify mechanisms and predictors of heterogenous treatment responses.

#### The social dimension of psychedelic therapy

Psychiatric disorders such as depression, anxiety, PTSD, and addiction are all essentially social in their manifestation and impact. They disrupt social relationships, are exacerbated by social stress and isolation (120, 121), and often improve with social support and connection (122). Emerging evidence suggests that psychedelic therapy may exert therapeutic effects partly through improvements in social functioning: patients report enhanced feelings of connectedness, reduced social anxiety, increased empathy, and improved relationship quality following psychedelic experiences (123). Specifically, psilocybin reduces neural responses to social exclusion (124), enhances emotional empathy and prosocial behaviour (125, 126), improves social adaptation during interpersonal feedback (127), and increases feelings of trust and closeness to others (126). What’s more, one of the central predictors of psychedelic therapy outcomes is the social bond formed with the supporting therapist(s) (12, 13), further highlighting that the therapeutic benefits of psychedelics are essentially social in nature. These findings suggest that understanding psychedelic mechanisms requires studying not just isolated individuals, but individuals embedded within a social context — how they navigate social hierarchies, form affiliations, respond to social stress, and how their position within social networks shifts over time.

Despite the deeply social nature of both psychiatric disorders and psychedelic therapy, preclinical psychedelic research has largely studied individual animals in isolation. This approach overlooks the possibility that psychedelics may exert therapeutic effects partly by altering how individuals interact with others. Recent methodological advances make it feasible to address this gap: automated behavioural tracking can now characterize social interactions in groups of animals over extended periods (128–130), social network analysis can reveal how psychedelic treatment alters an individual’s position within a social hierarchy (131, 132), and wireless neural recording systems enable brain activity monitoring during naturalistic social behaviours (133, 134). Such approaches could reveal whether psychedelics alter social approach-avoidance behaviours, modify responses to social stress, change patterns of affiliative interaction, or shift an individual’s dominance status. Moreover, studying individuals within their social context may be essential for capturing clinically relevant predictors, such as differing social positions, or adherence to distinct behavioural phenotypes (e.g., stress-responsive versus stress-resilient) (92). Given that social and empathy deficits characterize treatment-resistant depression (121) and that improved connectedness mediates therapeutic outcomes (12, 135), animal models that fail to take social functioning into account may miss the very mechanisms most relevant to clinical translation.

#### Long-term persistence and temporal dynamics

One of the most remarkable features of psychedelic therapy is the persistence of therapeutic effects long after acute effects have subsided. In clinical trials, single or double dosing sessions produce symptom improvements that can last for months or even years (4, 5). This temporal profile — brief intervention, lasting change — implies that psychedelics rapidly initiate neurobiological processes that continue to unfold long after the drug has been metabolized.

Animal research has made substantial progress in identifying candidate mechanisms for these lasting effects, particularly focusing on neuroplasticity-related dynamics such as increased dendritic spine density, enhanced synaptic plasticity, and structural remodelling of cortical circuits (136–139). Some studies have achieved commendable longitudinal follow-ups, with the longest extending to approximately one month post-treatment (136, 140, 141)—a substantial commitment given the practical challenges of sustained monitoring in animal research.

Yet here lies a largely untapped opportunity unique to preclinical models: in humans, ethical and practical constraints limit our ability to study individuals for extended periods, whereas animal models allow us to track the same individuals across their entire lifespan—from early development through adulthood and into old age. This capacity for lifelong monitoring represents a profound advantage rather than merely a gap to fill. While human therapeutic effects persist for months or years, we can directly observe how psychedelic-induced changes unfold across developmental transitions, ageing processes, and shifting social contexts that would be impossible to capture in clinical trials. Moreover, behavioural assessments in animal studies typically occur at discrete time points rather than tracking phenotypic changes longitudinally in the same individuals (140, 142), and social behaviour dynamics remain largely uncharacterized despite their documented importance in human therapeutic outcomes (143).

Long-term longitudinal designs within a social context are essential to address this gap. Such approaches would allow us to identify early neural, behavioural, and social predictors of lasting change, to characterize the temporal evolution of both individual circuit reorganization and behavioural and social shifts beyond the initial days, and to link transient pharmacological effects to sustained therapeutic mechanisms. Moreover, by combining continuous behavioural tracking of individuals within social groups with repeated neural recordings, we could directly test mechanistic hypotheses about how acute circuit perturbations translate into the persistent changes in brain function, individual behaviour, and social functioning that characterize therapeutic response in humans.

#### Practical constraints and intermediate steps

The reorientation outlined above can pose real challenges in terms of its implementation. In this section, we list six common practical constraints, and sketch solutions that move toward a more naturalistic experimental setting without requiring alternative infrastructure to be fully in place at once.

#### Voluntary administration and dose–response

Voluntary oral administration historically traded the stress of injection for loss of bolus timing and per-animal dose precision. A solid vehicle (peanut butter, chocolate spread, gelatin) consumed within a pre-trained 5–10 minute window largely solves both: intake is bolus-like, and weighing the offered aliquot before and after gives a per-animal dose covariate. Dose–response curves can then be derived either within-subject across days or between-subject across cohorts. For compounds whose oral pharmacokinetics diverge from their clinical pharmacokinetics, a dose-equivalence analysis remains required.

#### Longitudinal identity tracking

Within-subject designs require keeping track of which animal is which across weeks, including periods when animals are housed in groups of indistinguishable conspecifics. Hybrid RFID + video footage can address this challenge: RFID chips are highly affordable, can be implanted with minimal discomfort (144), and provide a simple means to anchor animal identity continuously, while the video stream supplies the behavioural readouts. RFID thus resolves identity, and video resolves behaviour, with neither requiring the animals to be marked or separated.

#### Data scaling and analysis

Continuous, multi-animal recording over weeks generates large data volumes, but this is rarely a hard limit in practice. On-the-fly feature extraction — computing compact behavioural descriptors at acquisition time rather than archiving raw video, for example pose-estimation keypoints (e.g. DeepLabCut or SLEAP), tracked positions and zone-occupancy time series, or motion-energy summaries, while optionally retaining full-resolution clips only around triggered events — reduces a multi-gigabyte video stream to lightweight time series. This can be combined with sampled-epoch designs focused on events of interest (e.g. dosing and post-dosing epochs). Adopting a standardised storage format such as Neurodata Without Borders (NWB) further helps by packaging behavioural, video and neural streams together with consistent metadata in a single self-describing file, which keeps long-running datasets organised and makes them straightforward to share across labs and re-analyse with common tooling — a prerequisite for the cross-lab data sharing we advocate below. Beyond storage, the within-subject structure of these designs is itself a statistical asset. In a conventional group-mean comparison, differences between individual animals enter the analysis as residual noise that inflates the error term and erodes power. A within-subject mixed-effects design instead lets each animal act as its own baseline and models that between-animal variability explicitly as a random effect, so a nuisance source of variance becomes part of the model structure rather than unexplained error. The result is greater sensitivity for the same number of animals — particularly valuable for the heterogeneous, longitudinal designs we advocate, where repeated baseline-to-post-dose measurements within each animal are exactly the data that such models exploit.

#### Validating new behavioural readouts

The multidimensional readouts we advocate are pharmacologically under-characterised during psychedelics, even where they have been validated in adjacent contexts (97, 145). Closing this gap is a multi-track effort: every new readout should be paired with a classical anchor (HTR/WDS) in the same animals to build a bridge dataset; sensitivity and specificity should be established using a reference-compound panel (psilocybin, LSD, DOI, plus a non-hallucinogenic 5-HT_2*A*_ agonist such as lisuride as a negative control (146, 147)); raw video and feature streams should be shared across labs; and a dose–response against a reference compound is required before a new readout can be treated as a translational tool.

### Chronic neural recordings in social groups

Tethered head-stages often tangle in group cages, and the implant surgery and tethering procedures are themselves stressors. In rats, wireless head-stages now permit chronic group recordings. In mice, no comparable wireless system is yet routinely deployed for adult-group recordings (134); the pragmatic alternative is to keep housing and behavioural testing group-based while confining the neural recording to individual sessions. Surgical and tethering stress can be reduced by longer recovery windows, lighter implants, analgesia, and habituation to the rig (148). We would therefore currently recommend that designs requiring simultaneous recording from multiple freely interacting animals during dosing should be conducted in rats.

#### Foundations first?

A common counter-proposal to the - certainly effortful - modifications suggested above is that we can characterise basic circuit mechanism in controlled simple setups first, and layer in social and environmental complexity later. This sequencing relies on the assumption that circuit responses in the simple setup generalise to more complex ones. Our review suggests that this assumption is not entirely valid in psychedelic research: current “controlled” setups provide stress-confounded conditions, whose circuit-level effects may not generalise to the therapeutic conditions the field is trying to model. Translational research on depression and anxiety pursued exactly this strategy for decades (149, 150), and its translational record is widely acknowledged to be poor (151, 152). A growing body of work attributes this shortfall in part to the conditions in which the foundational work was done: the limited external validity of standardised, stress-confounded designs, and the chronic stress imposed by routine housing and testing, have both been directly implicated in the poor reliability and translation of preclinical psychiatric research (151, 153, 154). From this perspective, characterising the behavioural and contextual state during a psychedelic episode is the fundament that translational research on circuit-level mechanisms has to be anchored to. This concern is not specific to psychedelics: each welfare variable we audit above has independently been shown to modulate the response to non-psychedelic drugs — antidepressants, psychostimulants, opioids, and alcohol — and the rapid-acting plasticity drug ketamine shows a comparable environment-dependent reversal of effect (40, 155). We would expect psychedelics to be similarly sensitive, which suggests that welfare context is better treated as a general modulator of drug action than as a psychedelic-specific detail.

### Concluding Remarks

Our analysis identifies three interconnected domains where preclinical psychedelic research diverges from the conditions thought to shape the therapeutic success of psychedelics in humans, each representing both a current limitation and an opportunity for methodological advancement.

#### Set and setting

most animal studies employ stress-inducing conditions: non-enriched housing, inactive-phase testing, forceful handling, and injection-based administration. These approaches fundamentally diverge from the supportive, low-anxiety contexts that have been associated with positive therapeutic outcomes in human studies. Adopting some of the proposed welfare-oriented practices such as environmental enrichment, communal housing, gentle handling, and circadian-aligned testing would create experimental conditions more likely to engage the same neurobiological mechanisms underlying human psychedelic therapy.

#### Experiential dynamics

current behavioural assays typically capture isolated markers at discrete time points rather than the multidimensional, temporally evolving states that predict therapeutic outcomes. Advances in computational ethology now enable comprehensive behavioural characterization in environmentally complex settings, offering a path toward capturing the richness of psychedelic-induced experiential states.

#### Individual and social context

while treatment response heterogeneity and social functioning are central to psychedelic therapy, most animal studies use group-level analyses of isolated individuals, precluding investigation of individual trajectories and social dynamics.

Together, these considerations point toward a fundamental reorientation: from brief, cross-sectional testing of isolated animals toward longitudinal tracking of individuals within enriched social environments. Neuroscience now holds the tools to meet this challenge, from complex analysis of naturalistic behaviours (156–158) to chronic neuronal recordings (133, 148) to long-term tracking of group dynamics (111, 132, 159).

This new approach can offer unique insights alongside what we already know about psychedelics from classical studies: the capacity for lifelong monitoring across developmental stages impossible to achieve in human studies, as well as the connection of dynamics across the levels of receptors, systems and behaviour. Although we have framed these arguments around psychedelics, the underlying principle is not specific to them: welfare and behavioural context shape drug action broadly, and the poor translational record of stress-confounded models in depression and anxiety research is a cautionary precedent rather than a coincidence. By embracing these opportunities, preclinical research can move beyond demonstrating that psychedelics produce overall effects in animals towards understanding the neuronal, contextual, experiential, and individual factors that determine whether those effects translate into lasting therapeutic change.

## Acknowledgements

The authors thank Claudia Kernberger for assistance with figure design. This work was supported by the Max Planck Society.

## Data Availability

No new experimental data were generated for this review. The full corpus of papers screened, the per-paper consensus scores (B/E/D dimensions and housing/experimental conditions), the analysis code that produced the figures, and the exact prompts used by the LLM scoring pipeline are openly archived on Zenodo (https://doi.org/10.5281/zenodo.20662840; see Supplementary Methods) and developed at https://github.com/hummuscience/psychedelic-behavioral-review, with an interactive dashboard for exploring the data at https://hummuscience.github.io/psychedelic-behavioral-review.

## Author Contributions

**Muad Y. Abd El Hay**: Conceptualization, Methodology, Formal analysis, Investigation, Data curation, Writing – Original draft, Writing – Review & Editing, Visualization, Project administration. **Ana Cukić**: Conceptualization, Methodology, Investigation, Writing – Review & Editing, Supervision. **Marieke L. Schölvinck**: Methodology, Investigation, Visualization, Writing – Review & Editing, Supervision, Funding acquisition. **Martha N. Havenith**: Conceptualization, Methodology, Software, Investigation, Writing – Review & Editing, Supervision, Funding acquisition.

## Declaration of generative AI and AI-assisted technologies

During the preparation of this work the authors used Claude Opus 4.5 (Anthropic, San Francisco, CA, USA) in order to generate a general structure, reformulate sentences, and perform LaTeX edits to prepare the manuscript for journal guidelines. After using this tool, the authors reviewed and edited the content as needed and take full responsibility for the content of the published article.

## Materials and Methods

We comprehensively reviewed articles reporting the effects of psychedelics on rodent behaviour identified by a systematic PubMed search, restricted to publications from 1 January 2014 onwards, with a date cutoff of 1 April 2026. The following search prompt was used for the initial list of studies:

~~~
(psychedelics[title/abstract] OR psilocybin[title/abstract] OR
psilocin[title/abstract] OR
“lysergic acid diethylamide”[title/abstract] OR LSD[title/abstract] OR
DOI[title/abstract] OR
DMT[title/abstract] OR “N,N-dimethyltryptamine”[title/abstract] OR
“5-MeO-DMT”[title/abstract] OR
mescaline[title/abstract] OR
ibogaine[title/abstract] OR ayahuasca[title/abstract])
AND
(mouse[title/abstract] OR mice[title/abstract] OR
rat[title/abstract] OR rats[title/abstract])
~~~

This yielded 851 results, to which 11 further records identified from other sources were added, for a total of 862 records. An automated classifier removed 304 records that mentioned no psychedelic compound, leaving 558 records for title/abstract screening. At this stage we excluded a further 189 records: studies that were off-topic for a behavioural review (including reviews, human clinical trials, pharmacological studies without behavioural experiments, duplicates, and corrections; 92), studies of MDMA—which, despite often being grouped with psychedelics in clinical contexts, is pharmacologically distinct from classical 5-HT_2*A*_ agonist psychedelics (76)—and studies with no behaviour reported in the full text (21). Of the remaining 369 reports, 14 were inaccessible due to being behind a paywall, and a further 89 were excluded after full-text review because they reported no administered drug dose to live animals (81) or no scored behavioural assay (8), resulting in a total of 266 studies included in the review (Supplementary Fig. S1).

### Behavioural Assay Characterization

For each study, we extracted information about every behavioural assay used and scored it along three dimensions. **Behavioural complexity** captured the number of distinct measures, the presence of temporal dynamics, sequential structure, ethological relevance, social and cognitive components, and the use of multivariate analyses (eight items, maximum 12 per assay). **Environmental complexity** captured the apparatus type, shelter, bedding, nesting, enrichment, spatial layout, social context, and food/water availability of the testing environment (eight items, maximum 11). **Recording duration** captured continuity, session length, recording span, baseline assessment, follow-up, and circadian coverage (six items, maximum 11). The full rubric with per-item definitions and worked examples is provided as Supplementary Methods.

Scoring was performed by an LLM-assisted multi-rater pipeline: each paper’s full text was converted to markdown via Docling, scored independently by three large language models (Qwen3-VL, Z.AI GLM-4.6, OpenAI GPT-OSS-120B), and the three outputs were reconciled into a consensus score by a fourth model (Claude Opus 4.7) acting as a judge. The judge’s per-item outputs include an evidence quotation extracted verbatim from the paper, ensuring every score is traceable. To validate the pipeline, the consensus scores were compared against two independent human raters on the assays scored by all three (*n* = 41; Supplementary Fig. S7). The standard deviation of paired score differences was of comparable magnitude across all rater pairs, including human–human: for behavioural complexity it was 0.72 between the two hu-man raters versus 0.86 (rater 1) and 0.55 (rater 2) against the consensus; for environmental complexity 0.50 versus 0.75 and 0.69; and for recording duration 1.60 versus 1.74 and0.91. Exact-agreement rates followed the same pattern (e.g. behavioural complexity 66% between the two human raters versus 63% and 78% against the consensus). One human rater diverged from both the other human rater and the consensus on two recurring item types—the count of distinct behavioural measures and the number of recording days per assay—and spot-checking these cases against the source papers confirmed that such disagreements reflect genuine interpretive ambiguity in those items rather than a deficiency specific to the automated pipeline. Recording duration was the least reproducible dimension for all rater pairs, indicating intrinsic ambiguity in that dimension rather than a limitation specific to the automated scoring. For each dimension, the study-level score reported in Figure 2 is the maximum across the study’s assays.

## Supplementary Information

For each behavioural assay, three dimensions were scored: Behavioural complexity (B1–B8, eight items, raw maximum 12), Environmental complexity (E1–E8, eight items, raw maximum 11), and Recording duration (D1–D6, six items, raw maximum 11). Items were scored from the recording session itself (when behavioural data are actively collected), not from conditioning, training, housing, or drug-administration phases. Each item was scored as 0 with confidence “low” and evidence “not reported” when the relevant information was absent from the paper, rather than inferred from what is typical for the assay. Items definitionally absent from the assay (e.g. bedding in an elevated plus maze) were scored 0 with confidence “high” and evidence “definitionally absent”.

### Supplementary Methods: Scoring rubric

Behavioural complexity (B1–B8).
| Item | Question | Response categories | Points |
| --- | --- | --- | --- |
| B1 | Number of distinct behavioural measures reported. Each unique behavioural variable counts once; sub-scores of the same composite each count separately; a derived preference index counts as 1. | 1 / 2–3 / 4–6 / 7+ | 0 / 1 / 2 / 3 |
| B2 | Temporal dynamics analysed within a single recording session (time-course plots, habituation curves, minute-by-minute analysis). Between-session comparisons do not count. | No / Yes | 0 / 1 |
| B3 | Sequential or transitional structure (transitions, behavioural sequences, Markov chains, bout structure, microstructure). | No / Yes | 0 / 1 |
| B4 | Ethological behaviours from the species’ natural repertoire directly measured during the recording session (grooming patterns, social investigation, nesting, burrowing, USVs). Head-twitch / wet-dog shake, locomotor distance, zone preference time, and freezing do not count. | No / Yes | 0 / 1 |
| B5 | Social component in the recording session. None: no social construct measured. Social construct, individual testing: assay targets social cognition but the focal animal is tested alone. Restricted social interaction: a live conspecific is present behind a barrier. Free social interaction: conspecifics interact without barriers and interaction is quantified. | None / Construct, alone / Restricted / Free | 0 / 1 / 2 / 3 |
| B6 | Cognitive or learned component (fear conditioning, Morris water maze, novel object recognition, reversal learning, operant tasks, conditioned place preference). Innate-avoidance anxiety tests do not count. | No / Yes | 0 / 1 |
| B7 | Automated behavioural classification beyond position tracking. Pose estimation (DeepLabCut, SLEAP), behavioural-syllable methods (MoSeq, B-SOiD, SimBA), or supervised ML classifiers count. Position-tracking systems (EthoVision, ANY-maze, Med Associates, beam-break) do not count. | No / Yes | 0 / 1 |
| B8 | Multivariate behavioural analysis (PCA, UMAP/t-SNE, factor analysis, cluster analysis across behavioural features). Multiple separate univariate tests do not count. | No / Yes | 0 / 1 |

**Environmental complexity (E1–E8).**
| Item | Question | Response categories | Points |
| --- | --- | --- | --- |
| E1 | Type of testing environment. Standard apparatus: open field, EPM, T/Y-maze, operant chamber, FST cylinder, 3-chamber. Home cage: animal's own housing as test environment. Enriched arena: larger or modified setup with added features. Semi-naturalistic: Visible Burrow System, IntelliCage, large multi-compartment or outdoor enclosures. | Standard / Home cage / Enriched / Semi-naturalistic | 0 / 1 / 2 / 3 |
| E2 | Shelter or refuge available to the animal during the recording session (nest box, igloo, tunnel, dark chamber accessed by choice). EPM closed arms do not count. | No / Yes | 0 / 1 |
| E3 | Bedding or floor substrate present in the testing apparatus during recording. | No / Yes | 0 / 1 |
| E4 | Nesting material available in the testing apparatus during recording. | No / Yes | 0 / 1 |
| E5 | Enrichment objects present in the testing apparatus during recording (tunnels, running wheels, chew toys, wooden blocks). Novel objects used as the test stimulus (NOR) do not count. | No / Yes | 0 / 1 |
| E6 | Spatial complexity. Multiple physically distinct areas separated by walls, dividers, or level changes. Open fields, FST cylinders, and EPMs count as single open spaces. | Single open space / Multiple zones | 0 / 1 |
| E7 | Social context during the recording session. Alone: focal animal only. Behind barrier: stimulus animal separated by mesh, divider, or wire cup. Free: $\geq 2$ animals interact without physical separation. Earlier conditioning phases do not count. | Alone / Behind barrier / Free | 0 / 1 / 2 |
| E8 | Food and water accessible inside the testing apparatus during the recording session. Home-cage food/water during tests run in a separate apparatus does not count. | No / Yes | 0 / 1 |

**Recording duration (D1–D6).**
| Item | Question | Response categories | Points |
| --- | --- | --- | --- |
| D1 | Recording continuity. Single: one recording bout on one day. Multiple: $\geq 2$ separate recording sessions with reported data. Continuous: uninterrupted recording for $\geq 6$ hours. | Single / Multiple / Continuous | 0 / 1 / 2 |
| D2 | Longest single session duration (longest individual recording bout). | <15 min / 15–60 min / 61 min–6 h / >6 h | 0 / 1 / 2 / 3 |
| D3 | Number of distinct calendar days on which behavioural data were collected for this specific assay (not the total study duration; not other assays' days). | 1 / 2–7 / 8–14 / 15+ | 0 / 1 / 2 / 3 |
| D4 | Pre-drug baseline recorded in the same assay and environment, with data reported or used in analysis. Habituation without data collection does not count. | No / Yes | 0 / 1 |
| D5 | Post-acute follow-up: behavioural recording $\geq 24$ hours after the last drug administration. | No / Yes | 0 / 1 |
| D6 | Circadian-scale coverage: any single recording session spans $\geq 6$ hours. | No / Yes | 0 / 1 |

#### Study-level aggregation

For studies that include multiple assays, the study-level score for each dimension reported in Figure 2 is the maximum across the study’s assays. This convention ensures that a study is credited for its most ambitious behavioural readout rather than penalised for also running short standard tests alongside.

#### Code and data availability

The exact prompt provided to the candidate scorers and to the judge, the per-paper consensus JSONs, the human-rater agreement analysis, and the analysis code that generated the figures in this manuscript are openly archived on Zenodo (https://doi.org/10.5281/zenodo.20662840) and developed at https://github.com/hummuscience/psychedelic-behavioral-review.

---

**Supplementary Figure S1.**
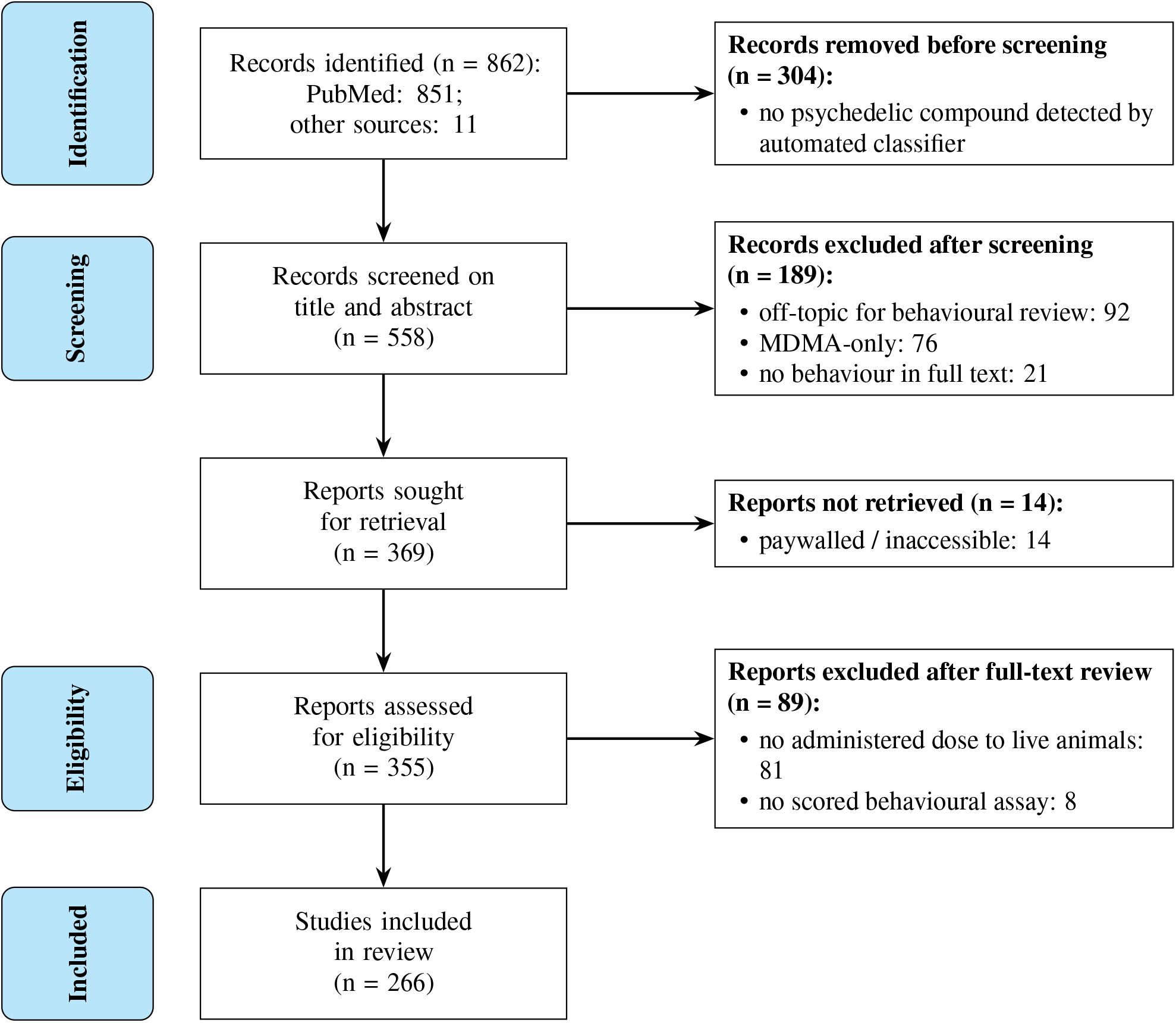
PRISMA 2020 flow diagram of the study selection process. Records were identified by an updated PubMed search of psychedelic-related literature (2014–2026), supplemented by a small number from other sources. An automated classifier removed records that mentioned no psychedelic compound prior to screening. Title/abstract screening further excluded off-topic, MDMA-only, and behaviourally empty records. After full-text review, the final corpus consists of studies with consensus LLM-extracted scores on at least one behavioural assay.

**Supplementary Figure S2.**
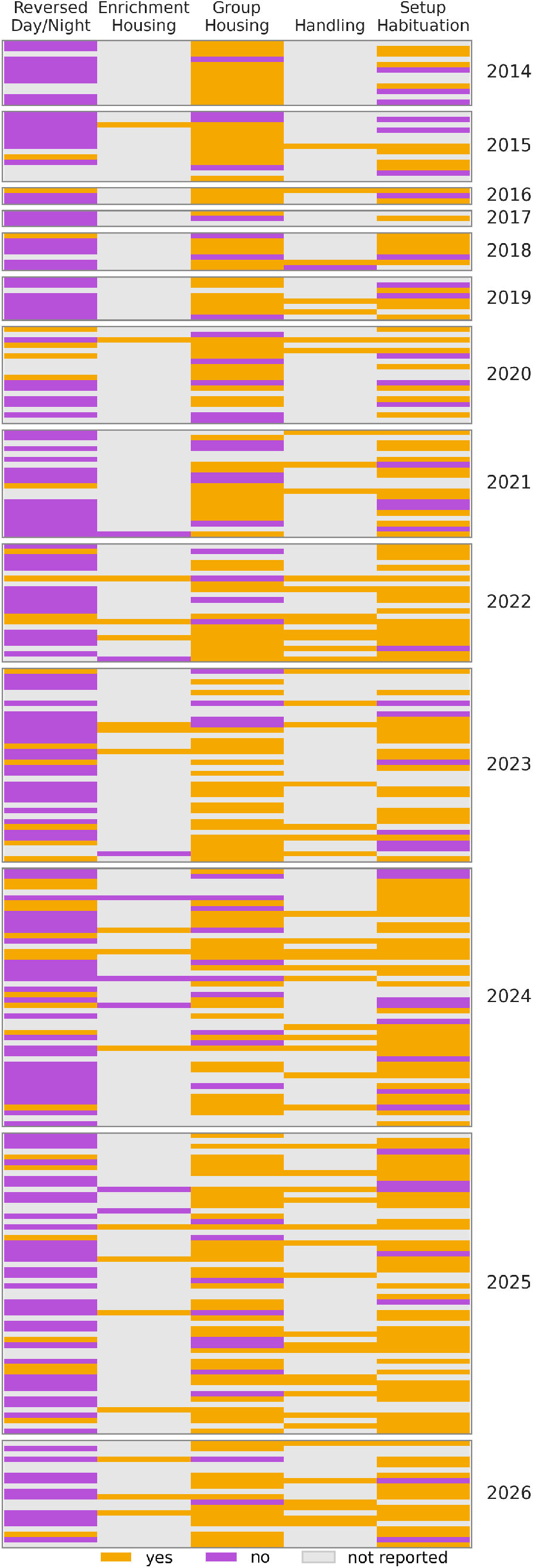
Systematic analysis of experimental conditions in psychedelic animal studies. Literature review for all studies using rodents in psychedelic research, from 2014–2026. Each row represents one study and each column one condition; cell colour indicates whether the practice was reported as used (orange), reported as not used (purple), or not reported (grey).

**Supplementary Figure S3.**
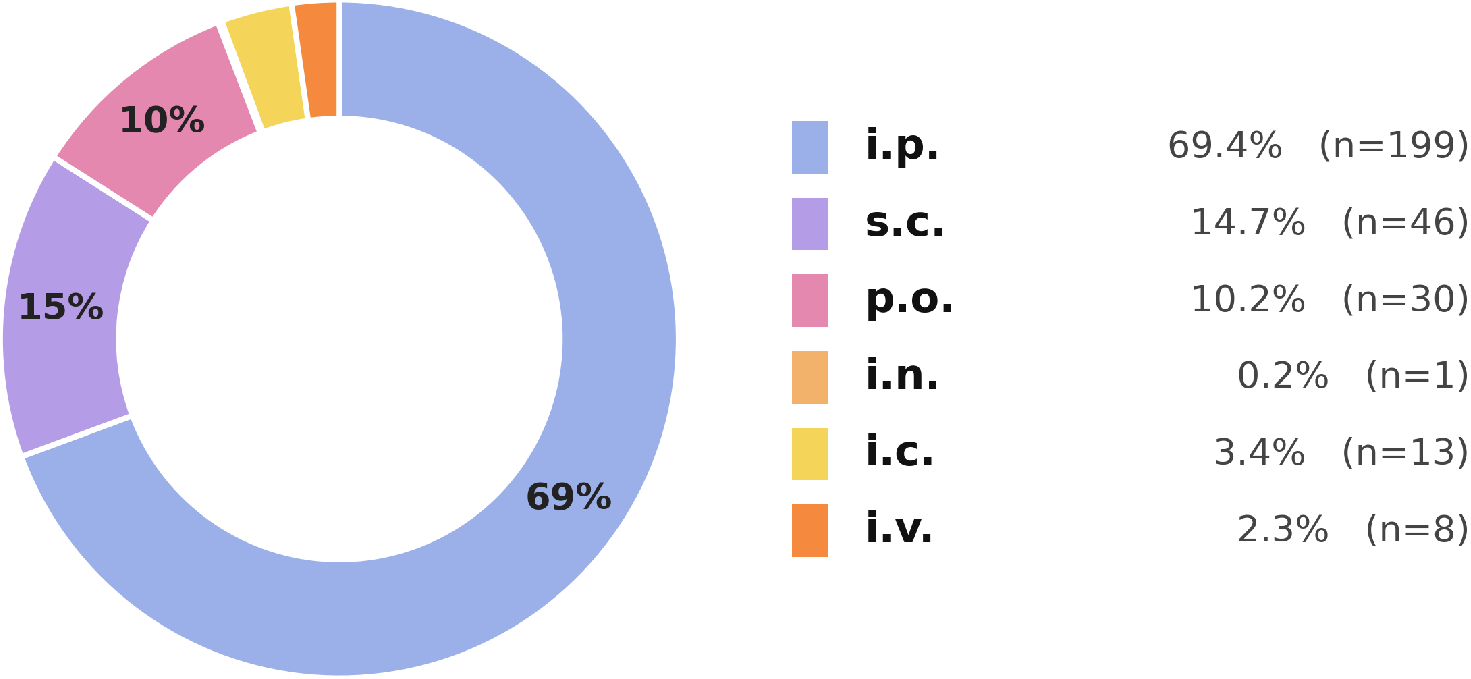
Distribution of psychedelic administration routes across the 266 reviewed studies. Each study contributes a total weight of one, split equally across the distinct routes by which its psychedelic(s) were administered, so that the shares sum to 100% over studies without double-counting a study that used more than one route. Intraperitoneal injection (i.p.) accounts for 69% of this weighted total (used in 199 studies), subcutaneous injection (s.c.) for 15% (46 studies), oral gavage (p.o.) for 10% (30 studies), intracranial microinjection (i.c.) for 3% (13 studies), intravenous (i.v.) for 2% (8 studies), and intranasal (i.n.) for less than 1% (1 study). Routes used only for non-psychedelic comparators or antagonists, and entries with no reported route, are excluded.

**Supplementary Figure S4.**
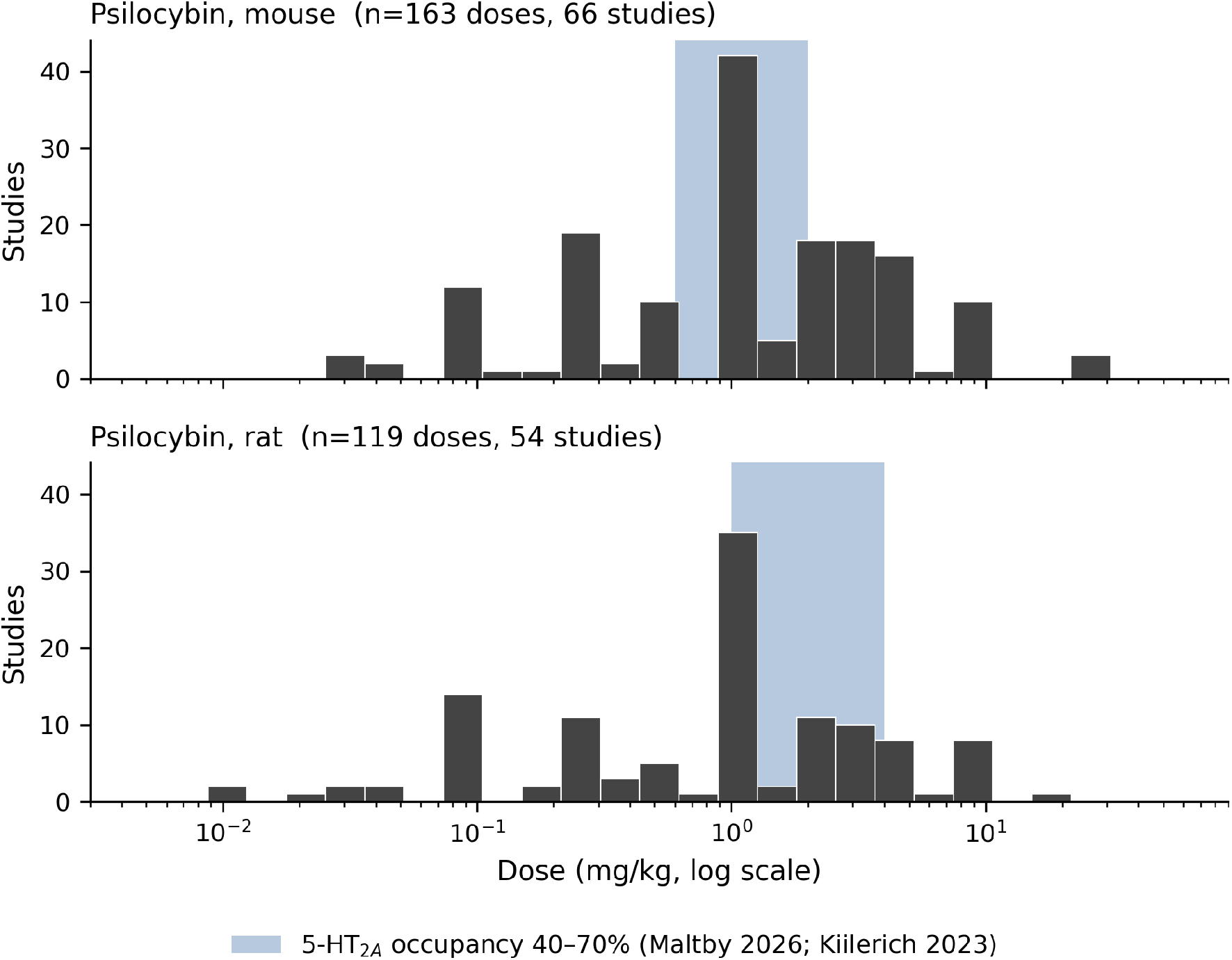
Distribution of psilocybin doses across the reviewed rodent studies, by species. Histograms of administered psilocybin (and psilocin) doses on a logarithmic scale for mouse studies (top) and rat studies (bottom). The shaded band marks the dose range corresponding to the clinically relevant 40–70% cortical 5-HT_2 *A*_ receptor occupancy window (mouse 0.6–2 mg/kg (78); rat 1–4 mg/kg (81)). The majority of reviewed doses in both species fall within or near this clinically equivalent range.

**Supplementary Figure S5.**
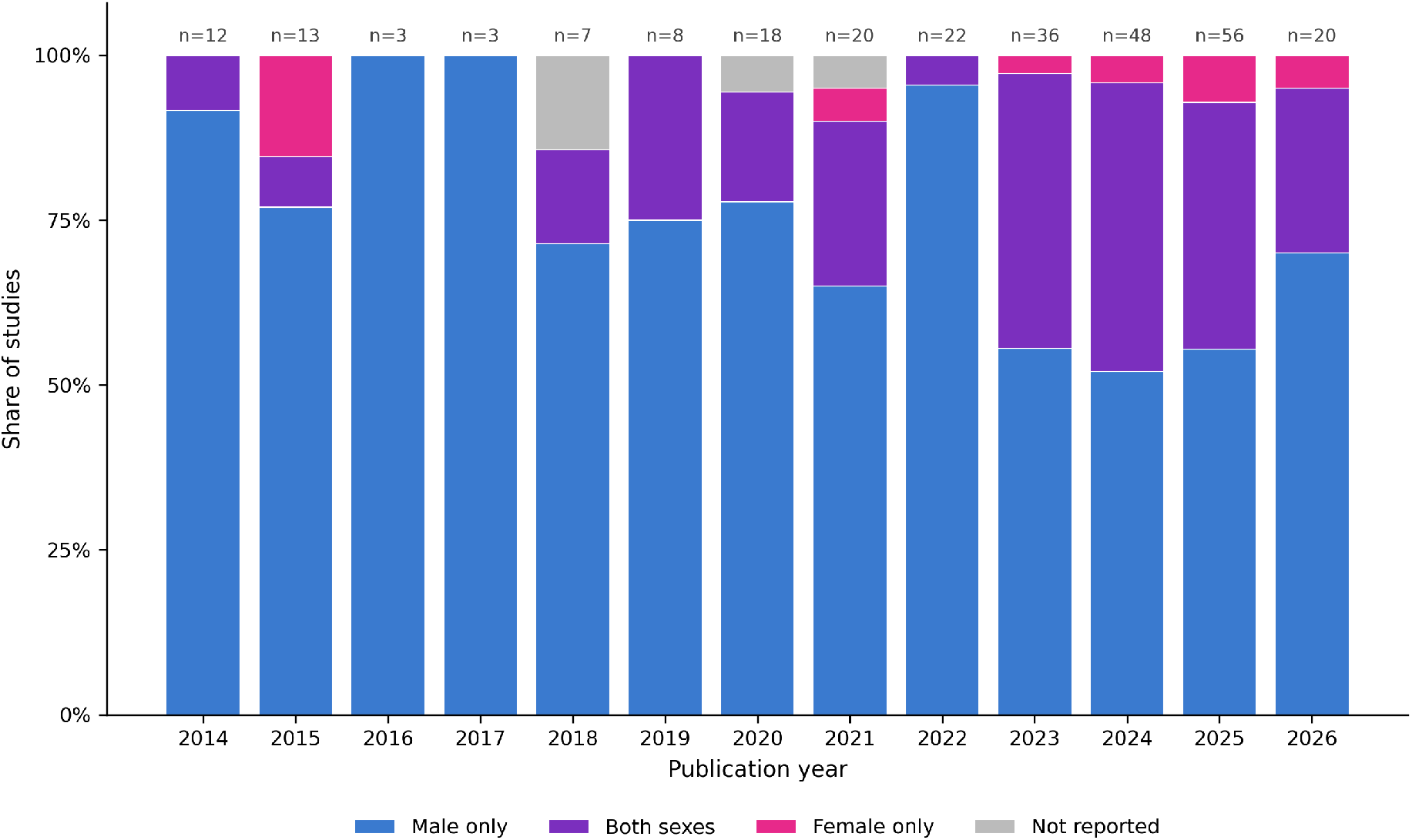
Sex composition of psychedelic rodent studies by publication year, 2014–2026. Each bar shows the share of studies in the corresponding year using male animals only, both sexes, female animals only, or not reporting sex. Total *n* per year is annotated above each bar. The share of studies including both sexes rose markedly from less than 15% in 2014–2021 to roughly 40% in 2023–2025, but male-only designs remained dominant throughout the period and female-only designs remained rare.

**Supplementary Figure S6.**
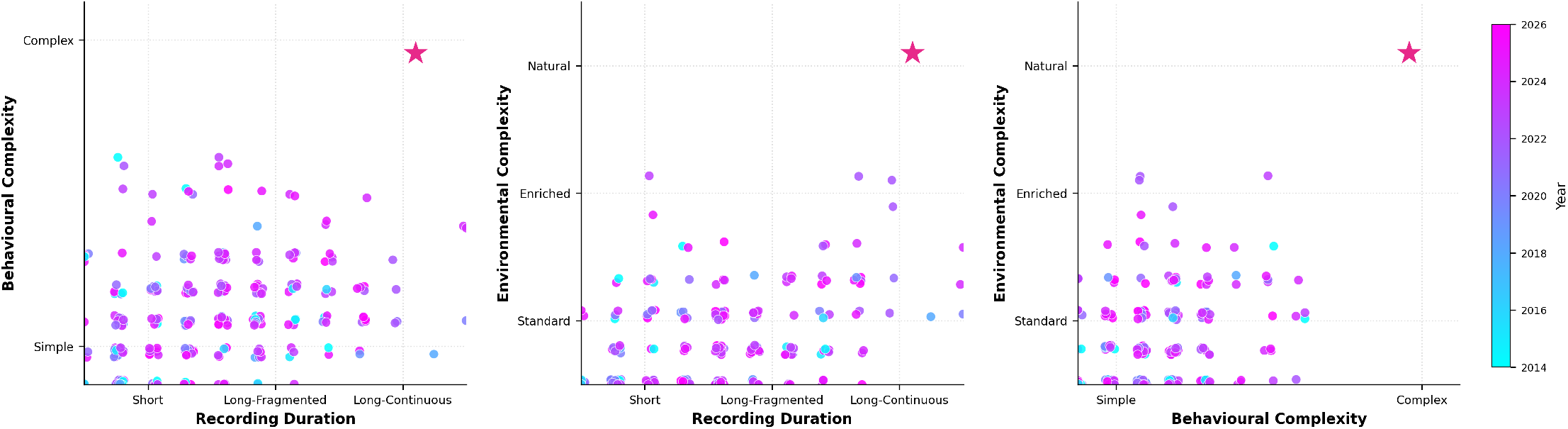
Two-dimensional projections of the behavioural-assay characterisation in Figure 2. The same 266 studies are plotted as three pairwise scatter plots: Recording Duration *×* Behavioural Complexity (left), Recording Duration *×* Environmental Complexity (middle), and Behavioural Complexity *×* Environmental Complexity (right). Each dot is one study, coloured by publication year. The pink star marks the aspirational target corner of each projection. Per-study positions are deterministically jittered by *±*0.2 so that overlapping integer-valued scores remain individually visible.

**Supplementary Figure S7.**
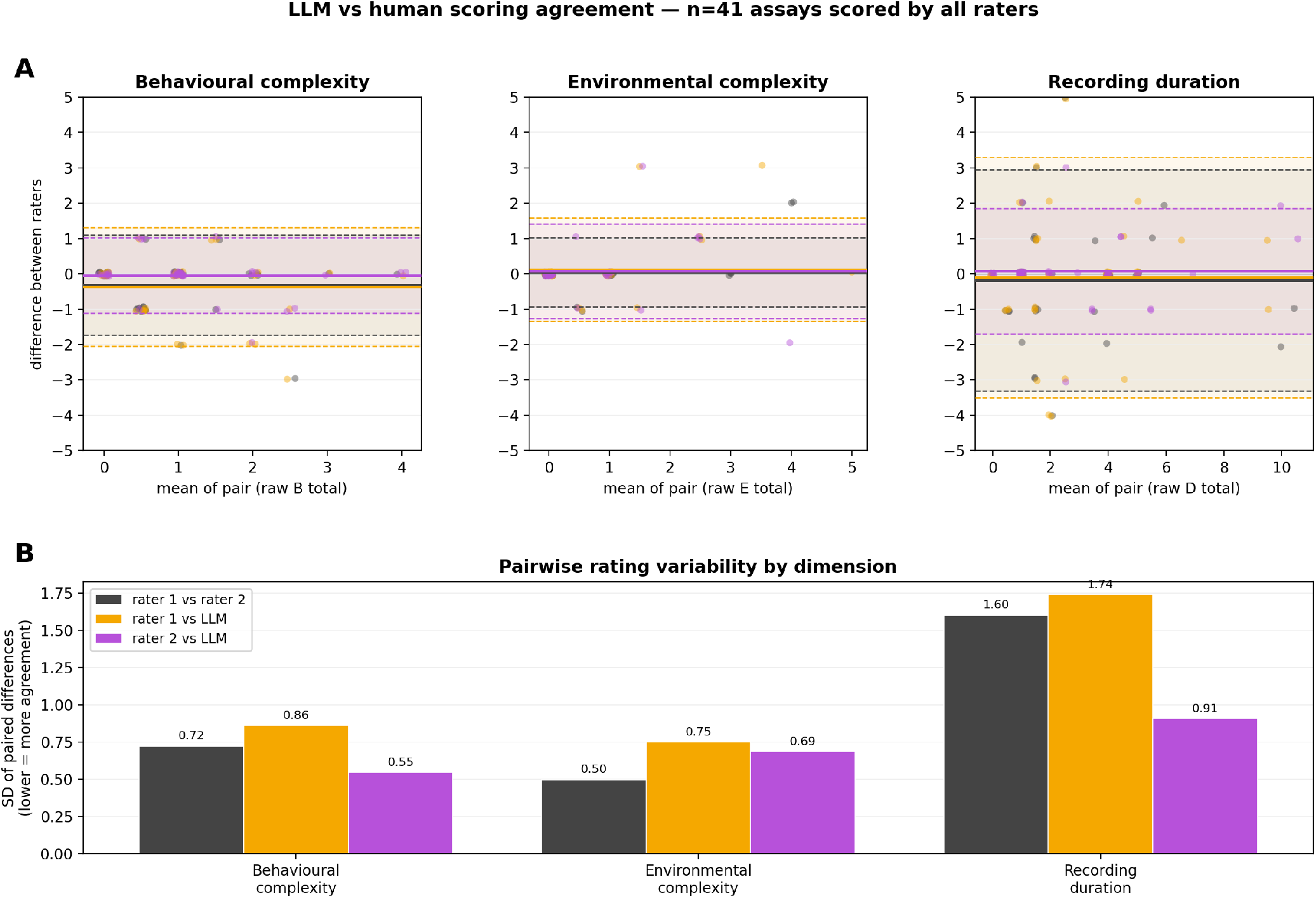
Inter-rater agreement on behavioural-assay scores. Agreement between three raters—two independent human raters (rater 1, rater 2) and the LLM consensus—across the *n* = 41 assays scored by all three, for the behavioural complexity, environmental complexity, and recording duration dimensions. **(A)** Bland–Altman plots per dimension with the three pairwise comparisons overlaid; solid lines indicate the mean difference (bias) and shaded bands the 95% limits of agreement. **(B)** Standard deviation of the paired score differences for each rater pair and dimension; lower values indicate closer agreement. *n* = 41 for behavioural and environmental complexity; *n* = 40 for recording duration.

**Supplementary Figure S8.**
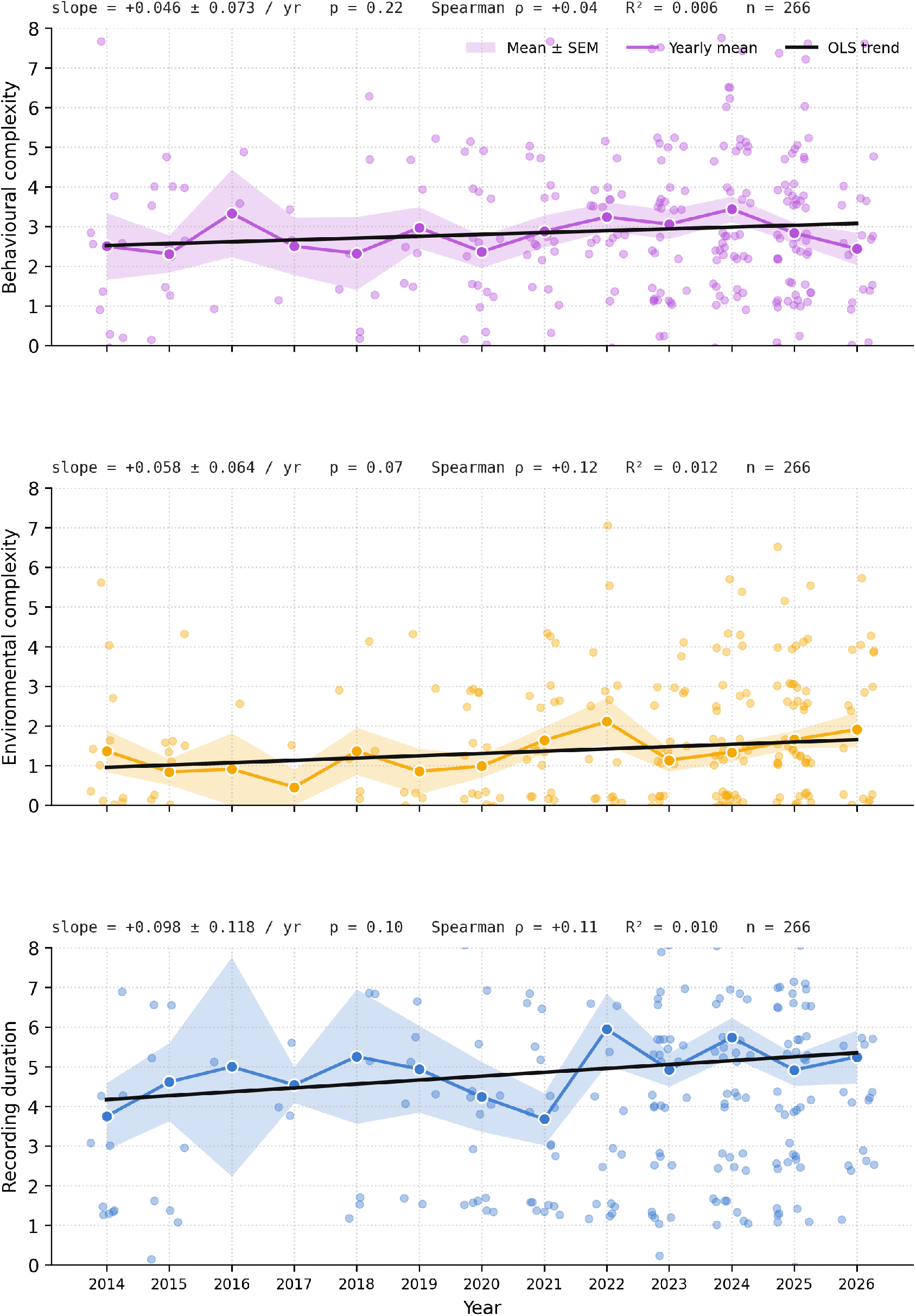
Temporal trends in behavioural-assay scoring dimensions, 2014–2026. Per-year scores for behavioural complexity (left), environmental complexity (middle), and recording duration (right) across the 266 reviewed studies. Solid lines show ordinary least squares fits with annotated slope estimates; shaded bands show the standard error of the mean. None of the three dimensions shows a significant temporal trend (behavioural slope +0.046 *±* 0.073/yr, *p* = 0.22; environmental +0.058 *±* 0.064/yr, *p* = 0.07; duration +0.098 *±* 0.118/yr, *p* = 0.10), indicating that the methodological reorientation we advocate has not been spontaneously emerging within the field over the past decade.

